# Macrophage-derived miR-21 drives overwhelming glycolytic and inflammatory response during sepsis via repression of the PGE_2_/IL-10 axis

**DOI:** 10.1101/2020.05.09.086421

**Authors:** Paulo Melo, Annie Rocio Pineros Alvarez, C. Henrique Serezani

**Author notes:** Equal contribution (Paulo Melo, Annie Rocio Pineros Alvarez). Vanderbilt University Medical Center. Division of Infectious Diseases, Department of Medicine. 1161 21^st^ Street South. A-220 MCN, Nashville, TN 37232-2582.

## Abstract

Myeloid cells play a critical role in the development of systemic inflammation and organ damage during sepsis. The mechanisms the development of aberrant inflammatory response remains to be elucidated. MicroRNAs are small non-coding RNAs that could prevent the expression of inflammatory molecules. Although the microRNA-21 (miR-21) is abundantly expressed in macrophages, the role of miR-21 in sepsis is controversial. Here we showed that miR-21 is upregulated in neutrophils and macrophages from septic mice. We found that myeloid-specific miR-21 deletion enhances animal survival, followed by decreased bacterial growth and organ damage during sepsis. Increased resistance against sepsis was associated with a reduction of aerobic glycolysis (as determined by reduced extracellular acidification rate (ECAR) and expression of glycolytic enzymes) and systemic inflammatory response (IL-1βTNFα and IL-6). While miR-21-/- macrophages failed to induce aerobic glycolysis and production of pro-inflammatory cytokines, we observed increased levels of the anti-inflammatory mediators’ prostaglandin E_2_ (PGE_2_) and IL10. Using blocking antibodies and pharmacological tools, we further discovered that increased survival and decreased systemic inflammation in miR21_Δmyel_ during sepsis is dependent on the PGE_2_/IL10-mediated glycolysis inhibition. Together, we are showing a heretofore unknown role of macrophage miR21 in the orchestrating the balance between anti-inflammatory mediators and metabolic reprogramming that drives cytokine storm and tissue damage during sepsis.

## Introduction

Sepsis is life-threatening organ dysfunction caused by a dysregulated host response to infection (1, 2). Regarding the last sepsis reports, 48.9 million incident cases of sepsis and 11 million sepsis-related deaths were reported worldwide in 2017 (3). The net between the infecting microorganisms and the host innate immune system culminates in a complex pathogenesis event associated with sepsis (4-6). Previous studies have demonstrated that the macrophage-mediated inflammatory response is involved in exaggerated systemic inflammatory and the immune suppression of sepsis (7-10). During sepsis, macrophages mediate the inflammatory response, metabolic switch dependent, and are heavily involved in the inflammation and immune suppression during sepsis(10-12).

Aberrant activation of pathogen recognition receptor (PRRs) by both pathogen or host-derived molecular patterns, leads to the activation of a myriad of pro-inflammatory programs that are triggered by nuclear factor-kappa B (NF-κB), Janus kinases (JAKs) and signal transducers and activator of transcription (STATs) and hypoxia-induced factor 1α (HIF1α) (13-15). Activation of these pathways induces pro-inflammatory programs characterized by the production of IL-6, TNFα, IL-1β, nitric oxide (NO) and KC, CXCL2 and macrophage inflammatory protein 2 (MIP-2), which are crucial to controlling of bacterial infection (16-20). Furthermore, HIF-α-mediated expression of glycolytic enzymes and transports leads to aerobic glycolysis in macrophages (21, 22). Aerobic glycolysis is a critical metabolic pathway required for macrophage activation and pro-inflammatory activity (23-25). The blockage of glycolysis in macrophages showed a crucial role in the reduction of systemic inflammation leads to improvement of sepsis outcome in experimental models (10, 21, 26). Exaggerated production of inflammatory mediators and aerobic glycolysis are critical determinants of organ dysfunction during sepsis (27, 28).

Recent studies have shown that miRNA expression is associated with severe illness and suggested it as potential mediators during sepsis (29). microRNA (miRNAs) are small non-coding RNA molecules (18-to 23-nucleotides) that regulate gene expression by translational repression or posttranscriptional suppression (30, 31). miRNAs have an essential role in several biological processes, such as development, differentiation, cell survival, and inflammatory response (32). miRNAs regulate phagocyte cytokine and chemokine response, antimicrobial effector function, pathogen recognition, and tissue repair. The microRNA-21 (miR-21) is highly expressed in different immune cells, such as T/B lymphocytes, monocytes, macrophages, and dendritic cells (33). I*n vitro* studies showed that several inflammatory stimuli as LPS and cytokines as IL-6, TFG-β, and TNF-α induce miR21 expression (34, 35). Our laboratory showed that miR21 deficiency accounted for the homeostatic generation of M2 macrophages by controlling the expression of STAT3 in macrophages (36). On the other hand, miR21 can prevent phagocytosis (by targeting GTPases) (37), TLR signaling (inhibiting MyD88 and IRAK), and suppress NFκB signaling(33). Therefore, in macrophages, miR21 works as a double-edged sword by regulating pro- and anti-inflammatory programs.

Experimental and clinical studies showed that miR-21 is upregulated in acute and sustained in late sepsis (38). However, the role of miR21 in LPS-induced endotoxemia or sepsis is inconclusive. McClure *et al*., have shown that treatment of BALB/c mice with a miR21 inhibitor enhanced animal survival and decreased bacterial loads after CLP (39). Whole-body miR21_-/-_ mice were also protected from septic shock by reducing inflammasome activation and IL-1β processing and secretion (40). Delivery of a miR21 antagomir was cardioprotective in septic mice by increasing the numbers of myeloid suppressor cells (38).

On the other hand, Barnett et al. did not identify any differences in animal survival when global miR21 deficient mice were CLP-induced sepsis (41). Although interesting, these manuscripts did not reveal the cellular and molecular targets by which miR21_-/-_ improves animal survival, and the role of miR21 in aberrant and damaging metabolism during sepsis remains completely unresolved. Here we hypothesize is that during increased miR21 expression in myeloid cells inhibits the PGE_2_/IL10 axis and drives aberrant glycolysis and organ injury and lethality during sepsis. These tangled modifications account for the decreased inflammatory response, lung injury, and animal mortality.

## METHODS

### Mice

Female and male C57BL/6 and LysM_cre_ mice (8 weeks old; weight, 18 to 23 g) were obtained from The Jackson Laboratory. miR-21_fl/fl_ mouse strain was generated by Dr. Ivan Mircea trough Ozgene Ptv Ltd (Perth, Australia). The miR-21_fl/fl_ and LysM_cre_ mice were initially crossed at Indiana University School of Medicine to generate miR21_Δmyel_ mice and then maintained at Vanderbilt University Medical Center. Mice were maintained according to National Institutes of Health (NIH) guidelines for the use of experimental animals with the approval of the Indiana University School of Medicine and Vanderbilt University Medical Center Committee for the Use and Care of Animals.

### Polymicrobial sepsis and endotoxemia induction

Sepsis was induced by Cecal ligation and puncture (CLP), as previously described (42). A 21 G needle was used to induce moderate sepsis. The survival rate was evaluated 6 h after CLP and twice per day every day during 7 days after CLP induction. Peritoneal lavage, blood, and lung were harvested 6 h after surgery to measure cytokines, markers of tissue damage, or bacterial burden in septic mice.

In another experimental setting, miR21_fl/fl_ and miR21_Δmyel_ mice were challenged i.p. with either a moderate (5mg/kg) or lethal dose (10 mg/kg) of LPS from *E. coli* serotype 0111: B4 (Sigma-Aldrich) or 0.9% saline control (43). Anti-IL10R treatment or isotype control (10mg/kg, clone 1B1.3A BioXcel) was injected 1 hour before LPS injection (44). Mice survival was assessed over 7 days. Cytokine levels were measured in serum and/or peritoneal lavage fluid harvested 12 hours after challenge with LPS.

### Bacterial determination

Bacterial counts were determined as previously described (42). In brief, peritoneal lavage and blood were collected from septic mice 6 h after CLP surgery, and serial dilutions were plated on Muller-Hinton agar dishes (Difco Laboratories, Detroit, Michigan). Colonies were counted after incubation overnight at 37°C. The results are expressed as log CFU (colony-forming unit) per ml of blood or peritoneal exudate.

### Flow cytometry

Peritoneal cells were resuspended in FACS Buffer (PBS 1x, 2mM EDTA, 0,5% BSA). Cells suspension was blocked with CD16/CD32 (clone 2.4G2; BD Biosciences Pharmingen) for 10 min at 4C. The cells were stained with mouse anti-CD11b FITC (1:200, BD biosciences) for 30 min at 4C. BD Cytofix/Cytoperm™ was used to fixation and permeabilization of the cells according to the manufactory instructions (Biosciences Pharmingen). Antibodies used to detect proteins involved in glycolysis (Hk1 clone C35C4, Glut1 clone D3J3A and HIF-1a D1S7W) were from Cell Signaling (all at 1:500) were used to intracellular staining, followed by conjugate secondary ab (PE anti-IgG H+L 1:1000, Invitrogen). The cells were acquired by LSRII (BD Biosciences) at the VUMC Flow Cytometry Shared Resource core (FCSR). Data were analyzed with FlowJo Version 10.

### Determination of neutrophil migration

Mice were submitted to CLP and were sacrificed 6 h after CLP surgery. Cells present in the peritoneal cavity were harvested with PBS/EDTA solution. The cells were washed, depleted of red blood cells by hypotonic lysis, and quantified using a Neubauer chamber. Aliquots of cells (1×10_6_ cells /100 μL) were incubated with anti-FcγR antibodies to prevent nonspecific antibody binding, followed by incubation with phycoerythrin-conjugated CD11b mAb (BD Biosciences) and allophycocyanin-conjugated anti-Ly6G mAb (BD Biosciences). After incubation, the cells were washed, fixed, and analyzed by flow cytometry, using a BD LSR II flow cytometer.

### Cell isolation

Peritoneal cells from septic mice were harvest 12 hours after CLP. Ly6G_+_ cells were isolated by Neutrophil isolation Kit mouse and F4/80_+_ cells by Anti-F4/80 Microbeads Ultrapure mouse kit followed the manufactory instructions (MACS, Miltenyi Biotec).

### BMDM cell culture

Bone marrow cells were flushed from femurs and tibias of mice with 10mL of DMEM supplemented with 10% FBS (Gibco), antibiotic/antimycotic solution (100X, HyClone), GlutaMAX (100X, Gibco) and HEPES (25mM, Sigma). Cells were pelleted by centrifugation and resuspended in ACK lysis buffer for 5 min and resuspended in DMEM. 10_7_ cells were resuspended complete DMEM supplemented with 20 ng/mL of M-CSF (Peprotech) and 5 ng/mL of GM-CSF (Peprotech) and plated in macrophage plates. Three days later, more than 10 mL of BMDM medium was added to the culture. On day 6, the medium was replaced with 10mL of fresh BMDM medium, and 24 hours later, cells were treated with different doses of LPS from *E. coli* serotype 0111: B4 (Sigma-Aldrich) for different time points. Also, cells were treated with anti-IL10R (10 μg, clone 1B1.3A BioXcel), Cyclooxygenase inhibitors (Aspirin 100 μM, and Indomethacin 10 μM, Sigma Aldrich) and mPGES1 inhibitor CAY10526 (1 Ozgene Ptv M, Cayman Chem.).

### Cytokine and PGE_2_ determination

Cytokine and chemokine production from blood, peritoneal exudate, and homogenized lung were measured by ELISA according to the manufacturer’s instructions (Biolegend or R&D Systems). The levels of PGE_2_ were measured at the serum or the peritoneal lavage fluid of septic mice, as well in the BMDM supernatant. The ELISA was conducted according to the manufacture’s instruction (Cayman Chem.).

### Lactate activity assay

Lactate in peritoneal exudate was quantified using a Lactate Kit (Quibasa Quimica Basica - Brazil), according to the manufacturer’s instructions.

### Myeloperoxidase (MPO) assay

Lungs were harvested from mice at 6 h after CLP surgery and processed as previously described (45).

### Creatine kinase-MB (CK-MB) and glutamic-oxaloacetic transaminase (TGO) activity

The levels of CK-MB and TGO were measured in the serum of septic mice, 6 h after surgery as a biochemical indicator of heart and liver injury, respectively. The determinations were made using a commercial colorimetric kit (Labtest, Brazil).

### Gene expression analysis

Cells from BMDM culture, the peritoneal cavity, and bronchoalveolar lavage were harvested, and total RNA extraction was performed using the miRNeasy Mini kit (Qiagen) according to the manufacturer’s instructions and as previously described (46, 47). Relative gene expression was calculated using the comparative threshold cycle (C_t_) and expressed relative to control or WT groups (ΔΔCt method). Primers for *Rnu6* and *miR-21* were purchased from Sigma Aldrich. Primers for *β-actin, Tnf, Il-1 β, Il-6, Il-10, Nos2, Hif-1α, Hk-1, Hk-2, Scl2a-1, Pkm, Ldha, Slc16a3* were purchase from Integrated DNA Technologies **(table 1)**.

**Table 1:** Primers list

| Primer | Assay ID | Ref. Seq. |
| --- | --- | --- |
| *Il1β* | Mm.PT.58.41616450 | NM_008361(1) |
| *Tnf* | Mm.PT.58.12575861 | NM_013693(1) |
| *IL6* | Mm.PT.58.10005566 | NM_031168(1) |
| *IL6ra* | Mm.PT.58.31166746 | NM_010559(1) |
| *IL10* | Mm.PT.58.13531087 | NM_010548(1) |
| *HIF1a* | Mm.PT.58.11211292 | NM_010431(1) |
| *Hk1* | Mm.PT.58.9947184 | NM_001146100(2) |
| *Hk2* | Mm.PT.58.32698746 | NM_013820(1) |
| *Slc2a1* | Mm.PT.58.7590689 | NM_011400(1) |
| *Pkm* | Mm.PT.58.6642152 | NM_001253883(2) |
| *Ldha* | Mm.PT.58.29860774 | NM_001136069(2) |
| *Slc16a3* | Mm.PT.58.13307682.gs | NM_001038653(3) |
| *Nos2* | Mm.PT.58.43705194 | NM_010927(1) |
| *Actb* | Mm.PT. 39a.22214843.g | NM_007393(1) |
| *miR21* | M_Mir21_1 | NR_029738 |
| *RUN6* | MIRCP00001 |  |

### RNA sequencing and data Analysis

RNA was isolated from BMDMs as above and sent to VANderbilt Technologies for Advanced GEnomics (VANTAGE) core at Vanderbilt University. cDNA synthesis, end-repair, A-base addition, and ligation of the Illumina indexed adapters were performed according to the TruSeq RNA Sample Prep Kit v2 (Illumina, San Diego, CA). The concentration and size distribution of the completed libraries was determined using an Agilent Bioanalyzer DNA 1000 chip (Santa Clara, CA) and Qubit fluorometry (Invitrogen, Carlsbad, CA). Paired-end libraries were sequenced on an Illumina HiSeq 4000 following Illumina’s standard protocol using the Illumina cBot and HiSeq 3000/4000 PE Cluster Kit. Base-calling was performed using Illumina’s RTA software (version 2.5.2). Paired-end RNA-seq reads were aligned to the mouse reference genome (GRCm38/mm10) using RNA-seq spliced read mapper Tophat2 (v2.1.1). Each sample was analyzed in triplicate.

Raw read quality was assessed using FastQC (v0.11.5). Salmon (v0.14.0) was used to quantify transcript expression using annotation gencode v21 mouse transcriptome (https://www.gencodegenes.org/mouse/release_M21.html) under quant mode with default parameters (transcript index was generated with k = 31 using transcripts fa file from gencode v21). Transcript expression was summarized at the gene level and imported into R using txImport (48, 49). R package EnsDb.Mmusculus.v79 (https://bioconductor.org/packages/release/data/annotation/html/EnsDb.Mmusculus.v79.html) was used to annotate the genes. Then the differentially expressed genes were called using edgeR (v2.26.5) with Benjamini-Hochberg adjusted p-value < 0.05 and log2FoldChange > 2. R package clusterProfiler (v3.12.0) was used for the gene set over-representation analysis with KEGG database. K-means cluster was applied to genes expression values (normalized by count per million) using R function kmeans (from R package stats), the number of clusters was determined by total within sum of square. R package ggplot2 was used to make the volcano plots. Heatmap was generated using pheatmap (v1.0.12).

### Immunoblotting

Western blots were performed as previously described (47). Protein samples were resolved by Mini-PROTEAN_®_ TGX_™_ (Bio-Rad), transferred to a nitrocellulose membrane and probed with glycolysis antibody Sampler (Kit 8337, Cell Signaling) and β-actin Dylight 800 (1:1000, Invitrogen). As a secondary Abs, we used Donkey Anti-Rabbit IRDye_®_ 700/800 (LI-COR). The images were acquired in the LI-COR odyssey and the densitometry analysis performed by Image-J.

### Elicited macrophages

Peritoneal cavity from miR21_fl/fl_ or miR-21_Δmyel_ mice were elicited with injection of 1 ml 3% thioglycolate for four days, and elicited macrophages were collected and isolated as previously described (36). Phagocytes were stimulated with 10 ng/ml LPS (Sigma-Aldrich), and for indicated time points, the supernatant was used to determine cytokines production.

### Extracellular Flux Analyses (Seahorse)

Glycolytic stress assay (Extracellular acidification rate - ECAR) was performed using the Seahorse Bioscience XFe96 Extracellular Flux Analyzer. BMDMs from miR21_fl/fl_ or miR21_Δmyel_ in a density of 50,000 cells/well were used for all assays. BMDMs were seeded in a Seahorse 96 - well plate in DMEM and incubate overnight at 37C an atmosphere of 5% CO_2_. In the next day, BMDMs were unstimulated (US) or stimulated with LPS (1,10 and 100 ng/mL), anti-IL10R (clone 1B1.3A, BioXcel) and CAY10526 (Cayman CC) for 12 and 24h. After time points of stimulation, the culture medium was replaced for XF Base Media (DMEM) with 2mM glutamine supplemented. All experiments were done under sterile conditions and the freshly prepared media at a pH of 7.4 before each assay. The injection of glycolysis test compounds went on in this sequence: Glucose (Final concentration, FC 10 μM), Oligomycin (FC 1.0 μM), and 2-deoxyglucose (2-DG, FC of 50 μM). Glycolysis stress was measured by recording extracellular acidification rates (ECAR, milli-pH units per minute).

### Statistics analysis

Data are expressed as mean ± SEM obtained from two independent experiments. For comparisons between two groups was used test t, one-tailed for parametric data. Experiments with three or more groups were analyzed using one-way ANOVA and corrected using the Bonferroni multiple comparison test. Survival curves were expressed as percent survival, and Log-rank (Mantel-Cox) test was used to determine differences between survival curves. Results were analyzed with Prism 5.0 software (GraphPad Software, San Diego, CA). p < 0.05 was considered significant.

## RESULTS

### miR21 expression during sepsis

Initially, we evaluated the expression of miR-21 in mice undergoing polymicrobial sepsis induced by CLP-induced sepsis over time. Our data show that miR21 abundance is increased after 24 h and remains enhanced up to day 7 post-infection **(Figure 1A)**. Next, we aimed to assess whether macrophages or neutrophils express different levels of miR21 in both sham and septic mice. miR21 expression is higher in neutrophils than macrophages in naïve mice, but 24 h after sepsis, miR21 abundance was similar in both cells **(Figure 1B).** These data suggest that miR21 is abundantly expressed during sepsis in both macrophages and neutrophils.

**Figure 1:**
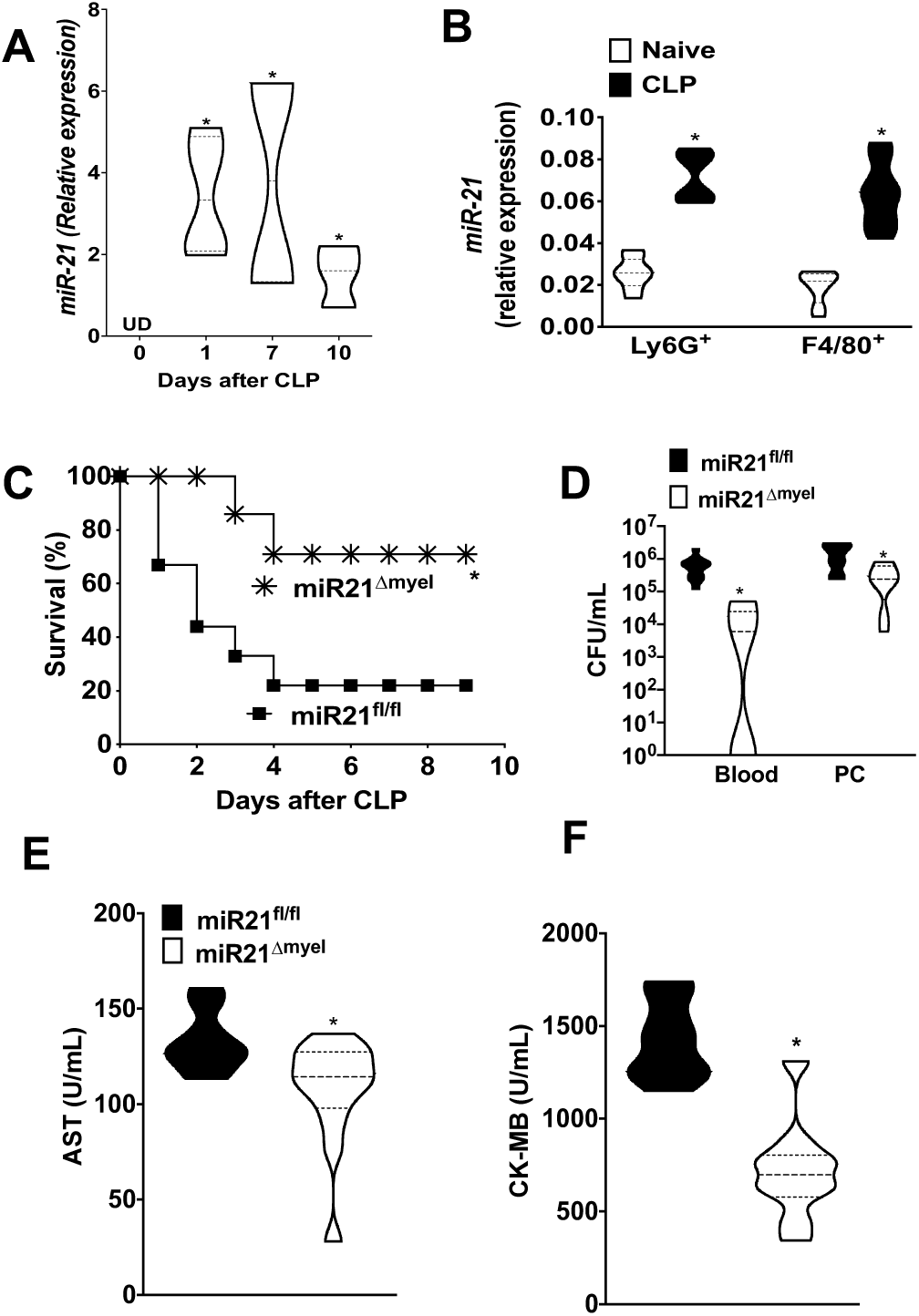
Myeloid-miR21 upregulation leads to poor sepsis outcomes. **(A)** miR-21 expression levels in the peritoneal cells from septic mice 1, 7, and 10 days after CLP determined by qPCR. **(B)** miR-21 expression in the Ly6G_+_ and F4/80_+_ peritoneal cells 12 hours after CLP by qPCR. Data are from *n* = 8 mice per group and were analyzed by *t-test* and Mann-Whitney *U* test. **(C)** The survival rate of miR21_fl/fl_ and miR21_Δmyel_ septic mice monitored for 7 days after CLP. Data are from *n* = 10 mice per group and were analyzed by the log-rank (Mantel-Cox) test. **(D)** Bacterial burden in the blood and peritoneal lavage of miR21_fl/fl_ and miR21_Δmyel_ determined by CFU counts. **(E**,**F)** AST (liver) and CKMB (heart) damage markers measured in the serum as described in the methods. **D-F** Data are from *n* = 7 mice per group and were analyzed by *t-test* and Mann-Whitney *U* test. Violin plots show the frequency distribution of the data. For the appropriate panels, \**P* < 0.05 compared to sham mice or miR21_fl/fl_ CLP (control) in all experiments.

### Absence of miR-21 in myeloid cells improves survival in septic mice

The role of miR21 during LPS-induced endotoxemia or sepsis is inconclusive. While McClure *et al*., have shown that miR21 antagomir enhanced animal survival and decreased bacterial loads after CLP in BALB/c mice (39). Enhanced survival was also associated with cardioprotective effects in miR21-antagomir-treated mice (38). Furthermore, whole body miR21_-/-_ mice are also protected from LPS-induced endotoxic shock, but not CLP (41). Although interesting, these manuscripts did not reveal the cellular and molecular targets by which miR21_-/-_ improves animal survival, and the role of miR21 in aberrant and damaging metabolism during sepsis remains completely unresolved. To define a role for myeloid-derived miR-21 in sepsis, we performed CLP in both miR21_fl/fl_ and miR21_Δmyel_. We found that while WT mice are susceptible to sepsis (80% mortality), miR21_Δmyel_ are protected (25% mortality) (**Figure 1C)**. These findings correlate with the reduced bacterial burden in blood and peritoneal cavity of miR21_Δmyel_ septic mice, when compared with septic WT control (**Figure 1D**). Next, we aimed to determine whether decreased cytokine production during sepsis correlates with lower organ damage. We observed that miR21_Δmyel_ showed reduced production of AST (aspartate transaminase) and creatine kinase-MB (CK-MB), a cardiac and hepatic marker of tissue damage, respectively (**Figure 1E, F**). Together these data show that myeloid-specific miR-21 expression is detrimental during sepsis and negatively influences animal mortality and bacterial load.

### miR21 regulation of cytokine production drives mortality during sepsis

Mice mortality during sepsis is controlled by early and exaggerated production of inflammatory cytokine, followed by impaired microbial clearance in later time points. (13, 50). Since we did not detect any apparent differences in antimicrobial effector functions (data not shown), we aimed to study the role of miR21 in inflammation-induced mortality in sepsis; we challenged miR21_fl/fl_ and miR21_Δmyel_ mice with either lethal or sublethal doses of LPS. Our data show that miR21 deficiency protected mice from LPS-induced septic shock **(Figure 2A)**. Next, we determine the production of systemic cytokines in LPS-challenged and CLP mice. Our data show that myeloid-specific miR21 deficiency leads to a robust decrease in the production of IL6, IL1β, and TNFα in the serum of both LPS and CLP septic mice **(Figure 2B, D).**

**Figure 2:**
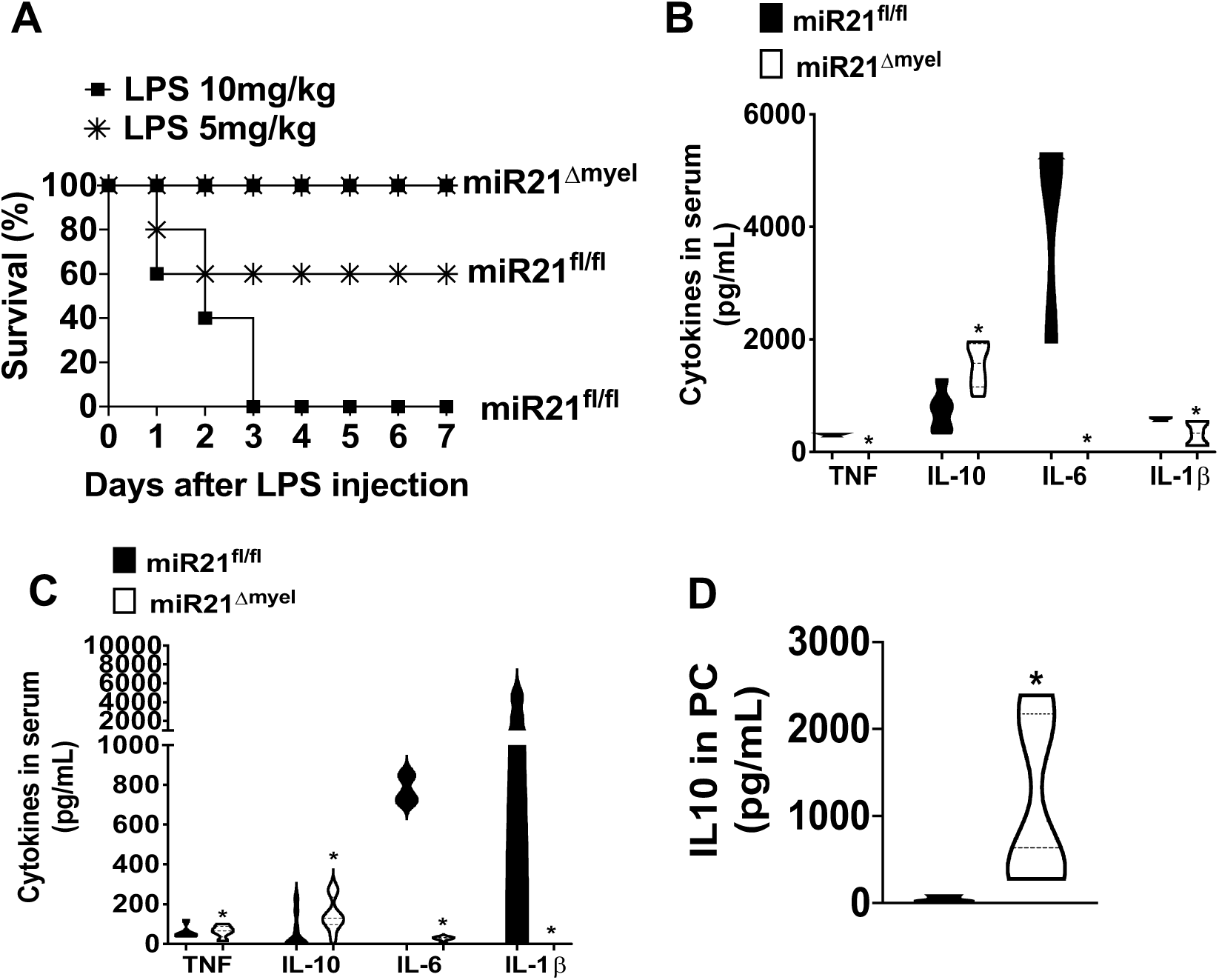
Myeloid-miR21 regulates cytokines levels in sepsis and septic shock. **(A)** Survival rate miR21_fl/fl_ and miR21_Δmyel_ mice challenged with lethal (10mg/kg) and sub-lethal (5mg/kg) doses of LPS. The survival curve was analyzed for 7 days (n=5/7 mice per group) \**P* < 0.05 versus LPS-treated miR21_fl/fl_ analyzed by log-rank (Mantel-Cox) test. **(B)** Concentrations of IL-1β, TNF-α, IL10, and IL-6 in the serum of miR21_fl/fl_ and miR21_Δmyel_ mice after 24h of a lethal dose of LPS injection by ELISA. **(C)** Concentrations of IL-1β, TNF-α, IL10, and IL-6 in the serum of miR21_fl/fl_ and miR21_Δmyel_ septic mice 18h after CLP by ELISA. **B, C** Data are from *n* = 4-7 mice per group and were analyzed by one-way ANOVA, followed by Bonferroni correction. **(D)** The concentration of IL-10 in the peritoneal exudate of miR21_fl/fl_ and miR21_Δmyel_ after 24h of LPS injection by ELISA. Data are from *n*= 4-7 mice per group and were analyzed by *t-test* and Mann-Whitney *U* test. Violin plots show the frequency distribution of the data. For the appropriate panels, \**P* < 0.05 compared to miR21_fl/fl_ septic mice in all experiments.

### miR-21 induces lung damage in septic mice

Recent studies have shown that miR-21 is increased in bronchoalveolar lavage fluid (BALF) cells, and it has a critical role in pulmonary inflammation (51). To further study the role of miR-21 in the lung of septic mice, initially, we evaluated miR-21 expression in cells from BALF. We demonstrated increased expression of miR-21 in the lung of septic mice, as compared to naïve mice **(Sup. Fig. 1A**). Next, we evaluated the migration of neutrophils to the lung of septic mice, and we found decreased lung-neutrophils sequestration and reduction of MPO activity and when compared to littermate control septic mice **(Sup. Fig. 1B, C**). Indeed, these findings correlate with decreased gene expression and production of IL-6 and IL-1β as compared to littermate control septic mice **(Sup. Fig.1D, F, G**). Together, these results show that miR-21 leads to an increase of pro-inflammatory cytokines production and infiltrating inflammatory cells in the lung during sepsis.

### miR21 is required for the global expression of genes involved in the inflammatory response

Giving the substantial and conflicting body of literature showing the effects of miR21 in macrophages, we sought to determine what are the potential candidates involved in improved survival and decreased inflammation in miR21_Δmyel_ using RNAseq analysis in both naïve and LPS-challenged BMDM from miR21fl/fl and miR21_Δmyel_. Our data clearly show a globally reduced gene expression in macrophages from miR21_Δmyel_ in both basal and 6 h after LPS challenge **(Figure 3A).** Next, we performed a heat-map analysis of genes expression in miR21_fl/fl_ and miR21_Δmyel_, and we observed a similar pattern as in **Figure 3A**. Although we observed a high number of inhibited genes in miR21_-/-_ macrophages **(Figure 3A and B)**, we observed an increase in essential gene clusters. We detected differential regulation of few gene clusters in naïve and LPS-challenged miR21_-/-_ BMDM. In naïve miR21 deficient cells, we observed a basal and homeostatic regulation of genes involved in the cell cycle, DNA replication, and oxidative phosphorylation (which could be involved in decreased glycolysis). When cells were challenged with LPS, we observed that genes that belong to cluster 6 are increased in both naïve and LPS challenged miR21_-/-_ cells. The relevant genes in this group are involved in PI3K (as expected), but also HIF1α TNFa signaling, antigen presentation and phagosome formation **(Figure. 3B)** Furthermore, pathway analysis of BMDMs challenged with LPS showed regulation of genes involved in metabolic programs and inflammatory cytokine generation **(Figure 3C)**. When we performed a heat-map analysis of the top 50 regulated genes, we observed a global decreased expression of critical genes involved in cell homeostasis, tissue homeostasis and repair and metabolisms, as well as inflammatory response, indicating that miR21 is required for overall cell homeostasis. We also observed a decreased expression of inflammatory and repair genes in miR21deficient macrophages. Overall, these data suggest that miR21 is a crucial mediator of macrophage homeostasis and a unique component of LPS responses and mortality during sepsis.

**Fig.3.**
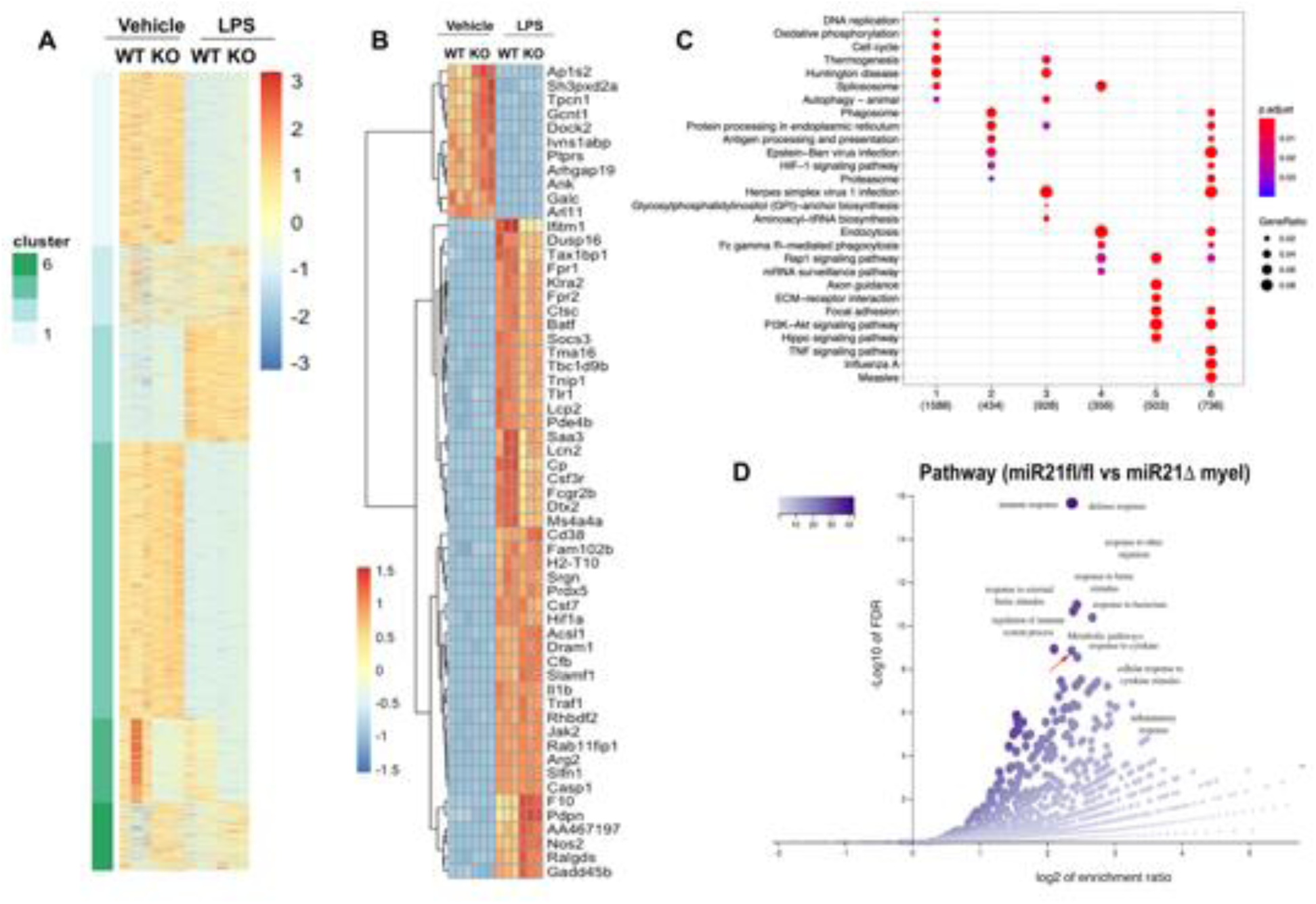
miR21 regulation of transcriptional programs in macrophages. BMDMs from miR21_fl/fl_ and miR21_Δmyel_ were challenged with LPS for 6 h, and RNAseq performed as described in the Methods. A) Heat-map of genes up-or down-regulated. Genes were clustered using a K-cluster tool (green bars on the right). B) Expression of the top 50 up or downregulated genes in. BMDMs as in A. C) KEGG analysis of gene clusters as in A. The X-axis shows the number of clusters and genes/clusters. D) Pathway analysis of genes from BMDMs as in A.

### miR-21 induces the production of pro-inflammatory cytokines in macrophages

Since our RNAseq data suggest an essential role for miR21 in the induction of inflammatory response, we aimed to investigate whether macrophages (the cells responsible for the systemic inflammatory response and organ damage during sepsis) (52, 53). We found that BMDM lacking miR-21 showed decreased IL-1β, IL6, and TNF-α mRNA expression (**Figure 4A, B, C**) when stimulated with LPS at different time points. Furthermore, peritoneal cells from septic mice also showed miR21 dependency for IL-1β, IL6, and TNF-α mRNA expression, but not for Nos2 mRNA expression **(Figure 4F).** Also, we confirmed the involvement of miR21 in the production of pro-inflammatory cytokines in response to LPS. We showed a reduction of IL-1β, IL6, and TNF-α in BMDMs, as well as a decrease of IL-6 production in peritoneal macrophages after LPS challenged **(Figure 4D, E, F, and Sup. Fig. 4).** However, the challenge of miR21_-/-_ BMDM with exogenous recombinant IL6 did not rescue LPS responses **(Sup. Fig. 5).** These data suggest that miR-21 drives the production of inflammatory cytokines in macrophages.

**Figure 4:**
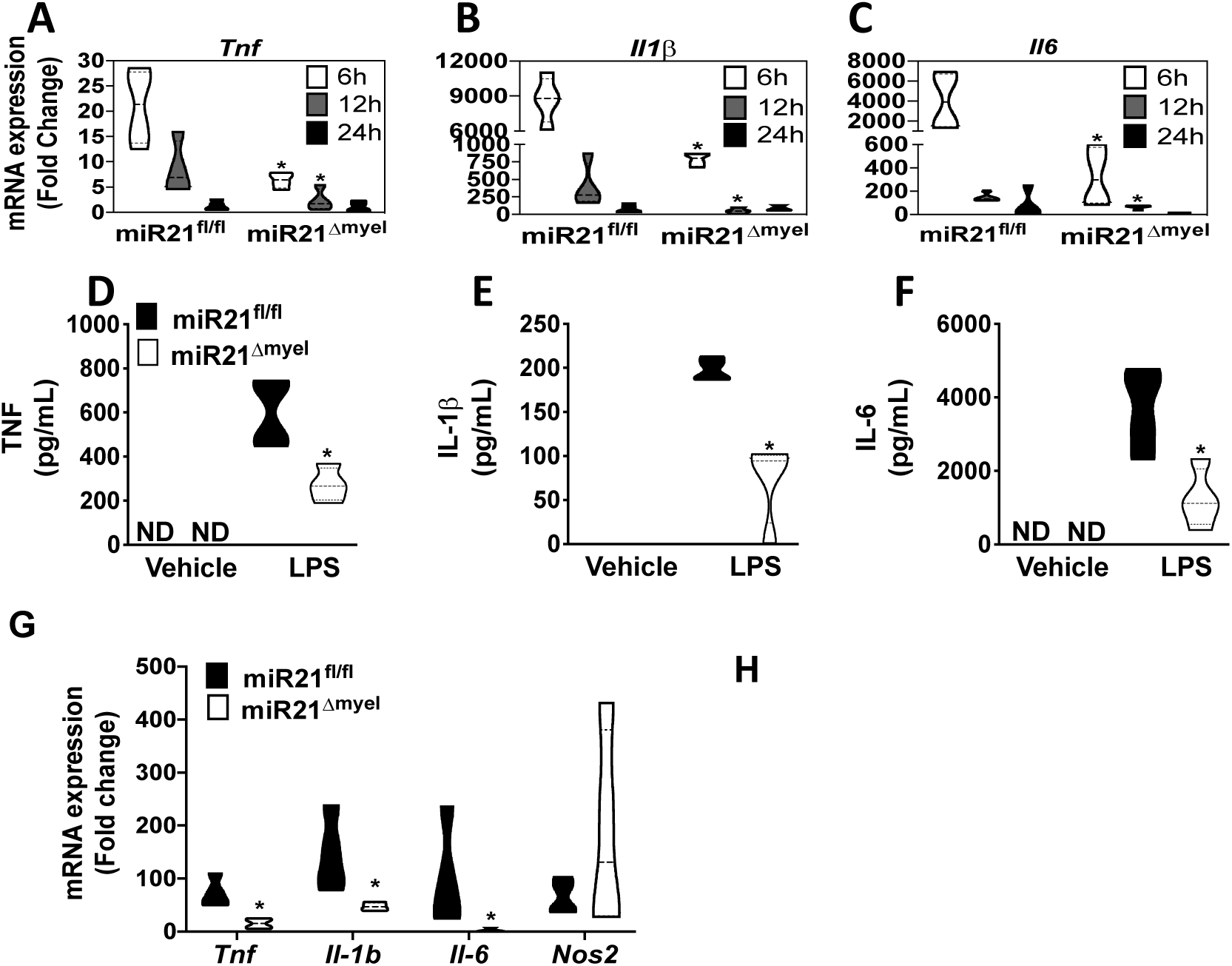
Expression of genes related to pro-inflammatory cytokines are miR21 dependent in myeloid cells. **(A-C)** Time course (6, 12, and 24h) of *Tnf, Il1b*, and *Il6* mRNA expression in miR21_fl/fl_ and miR21_Δmyel_ BMDMs challenged with LPS stimulation (10 ng/mL) determined by qPCR. **(D-F)** The concentration of TNF-α, IL-1β, and IL-6 in the supernatant of miR21_fl/fl_ and miR21_Δmyel_ BMDMs challenged or not with LPS (10ng/mL) for 24h determined by ELISA. For **A-F**, data are from *n* = 1-2 mice per group/experiment from at least 3 independent experiments and were analyzed by one-way ANOVA, followed by Bonferroni correction. **(G)** *Tnf, Il1b, Il6, and Nos2* mRNA expression in peritoneal cells 12h after CLP by qPCR. Data are from *n* = 4-7 mice per group and were analyzed by one-way ANOVA, followed by Bonferroni correction. Violin plots show the frequency distribution of the data \**P* < 0.05 compared in vivo by miR21_fl/fl_ mice or *in vitro* in LPS-challenged miR21_fl/fl_ BMDM (control) or unstimulated.

### miR21_-/-_ decreases glycolysis

Aberrant glycolysis drives the expression of inflammatory cytokines, enhances tissue injury and mortality during sepsis (10, 26, 54). Here, we speculated that miR21_-/-_ macrophages would exhibit lower glycolytic capability than WT cells challenged with LPS. Initially, we determined the extracellular acidification rate (ECAR) in macrophages from WT and *miR21*_*Δmye*l_ challenged or not with LPS using the Seahorse assay. Surprisingly, we detected lower basal glycolysis and glycolytic capacity (as measure by the ECAR) in LPS-challenged miR21 deficient cells than LPS-challenged WT, in both BMDMs and peritoneal macrophages **(Figure 5A, B, Sup. Fig. 4B and 4C).** These data led us to speculate that miR21 regulates the mRNA expression of enzymes involved in glycolysis. Peritoneal cells from *miR21*_*Δmye*l_ septic mice presented a downregulation of mRNA expression for most glycolytic enzymes (*Hk1, Pkm, and Ldha*), glucose, and lactate transporter (*Slc2a1 and Mct4*) compared to cells from WT septic **(Figure 5C).** Hk1 mRNA expression was downregulated in BALF cells from *miR21*_*Δmye*l_ septic mice **(Sup. Fig. 1E).** Moreover, BMDMs *miR21*_*Δmye*l_ LPS-challenged showed a significant reduction of glycolysis-related genes **(Sup. Fig. 4).** We also detected decreased protein expression of Hk1, GLUT1, and PDHK1 in miR21_-/-_ macrophages response to LPS in vitro **(Figure 5D)** and Hk1 and GLUT1 in CD11b_+_ peritoneal cells from *miR21*_*Δmye*l_ septic mice **(Figure 5E and Sup. Fig. 2A).** Next, we sought to determine whether the expression of key transcription factors involved in the expression of glycolytic enzymes was also controlled by miR21. Our data show that HIF1a expression is impaired in miR21_-/-_ cells ex vivo and in vitro **(Figure 5C, D, and E).** Since the ratio of pyruvate/lactate is decreased in cells undergoing aerobic glycolysis, leading to pyruvate-dependent lactate production (Ref). Next, we evaluated the abundance of pyruvate and lactate in peritoneal cells from septic mice 18h after CLP. We found lower levels of lactate and higher amounts of pyruvate in peritoneal cells from *miR21*_*Δmye*l_ septic mice when compared to WT septic animals **(Figure 5F, G).** Together, our findings show that miR21 deficiency protects mice from sepsis/endotoxemia by preventing glycolysis and lactate generation during sepsis.

**Figure 5:**
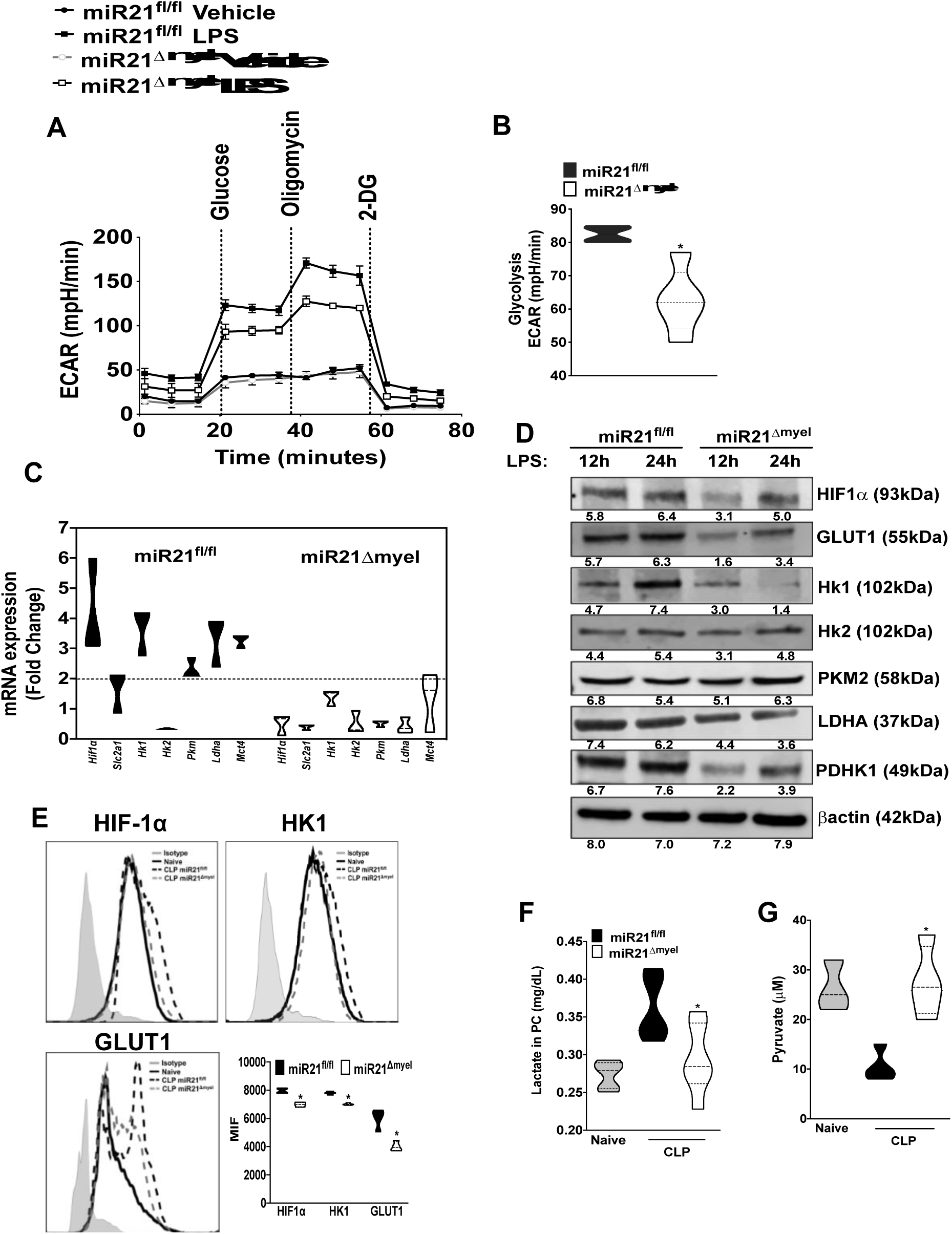
miR21 deficiency contributes to LPS-induced aerobic glycolysis resistance. **(A)** Real-time ECAR evaluation of glycolysis in miR21_fl/fl_ and miR21_Δmyel_ BMDMs after 12 hours of LPS (10 ng/mL) or PBS by Seahorse XF glycolysis stress test. **(B)** ECAR of glycolytic capacity after LPS. Data are from *n* = 1-2 mice per group/experiment from at least 3 independent experiments and were analyzed by one-way ANOVA, followed by Bonferroni correction. **(C)** *Hif1a, Scl2a1, Hk1, Hk2, Pkm, Lhda, and Slc16a3* mRNA expression determined in peritoneal cells from miR21_fl/fl_ and miR2_Δmye_ septic and sham mice 18h after surgery by qPCR. Data are from *n* = 4-7 mice per group and were analyzed by one-way ANOVA, followed by Bonferroni correction. **(D)** Expression of glycolytic proteins HIF-1 α, GLUT-1, HK1, HK2, PKM, PDHK1, LDHA, and βActin assayed in miR21_fl/fl_ and miR21_Δmyel_ BMDM after LPS-challenge of 12 and 24h by immunoblotting. Numbers underlanes represent densitometric quantification of two independent experiments. **(E)** Expression of HIF-1α, HK1, and GLUT1 in CD11b_+_ peritoneal cells from septic mice by flow cytometer. **(F-G)** Pyruvate and lactate measured in cells lysate and supernatant of the peritoneal cavity by colorimetric assay, respectively. Data are from *n* = 4-7 mice per group and were analyzed by one-way ANOVA, followed by Bonferroni correction. Violin plots show the frequency distribution of the data \**P* < 0.05 compared *in vivo* by miR21_fl/fl_ mice or *in vitro* by miR21_fl/fl_ BMDM LPS (control) or unstimulated.

### miR21 regulation of prostaglandin E_2_ production dictates the production of inflammatory mediators *in vivo* and *in vitro*

We and others have shown that PGE_2_ inhibits the production of pro-inflammatory mediators while enhances the production of anti-inflammatory actions of macrophages (36, 55, 56). Here, we speculated that PGE_2_ is a central mediator of the miR21 effects during sepsis/endotoxemia. Initially, we determined the abundance of PGE_2_ in the serum of miR21_flf_ and *miR21*_*Δmye*l_ mice. Our data show that decreased miR21 greatly enhances PGE_2_ during sepsis **(Figure 6A).** Next, we sought to investigate whether miR21 regulates basal and induced PGE_2_ levels in macrophages. Our data show that miR21_-/-_ BMDM produced PGE_2_ levels at both basal and after LPS challenge when compared to WT mice **(Figure 6B).** To determine whether increased basal PGE_2_ mediated low IL6 and TNFα and increased IL10, we treated BMDM from mir21_fl/fl_ and *miR21*_*Δmye*l_ with the specific mPGES-1 inhibitors CAY10526, followed by LPS challenge. Our data show that the mPGES2 inhibitor indeed decreases PGE_2_ abundance in *miR21*_*Δmye*l_ mice, and such effect is accompanied by increased pro-inflammatory cytokines production *miR21*_*Δmye*l_ BMDMs **(Figure 6C and D).** Next, we determined IL10 production in BDMD from miR21_fl/fl_ and *miR21*_*Δmye*l_ treated with different COX inhibitors (aspirin and Indomethacin, as well as the CAY10526 inhibitor). Our results clearly show that PGE_2_ inhibition decreases IL10 abundance in *miR21*_*Δmye*l_ macrophages **(Figure 6E).** Therefore, these data suggest that increased PGE_2_/IL10 leads to low pro-inflammatory cytokine production in LPS-treated cells.

**Figure 6:**
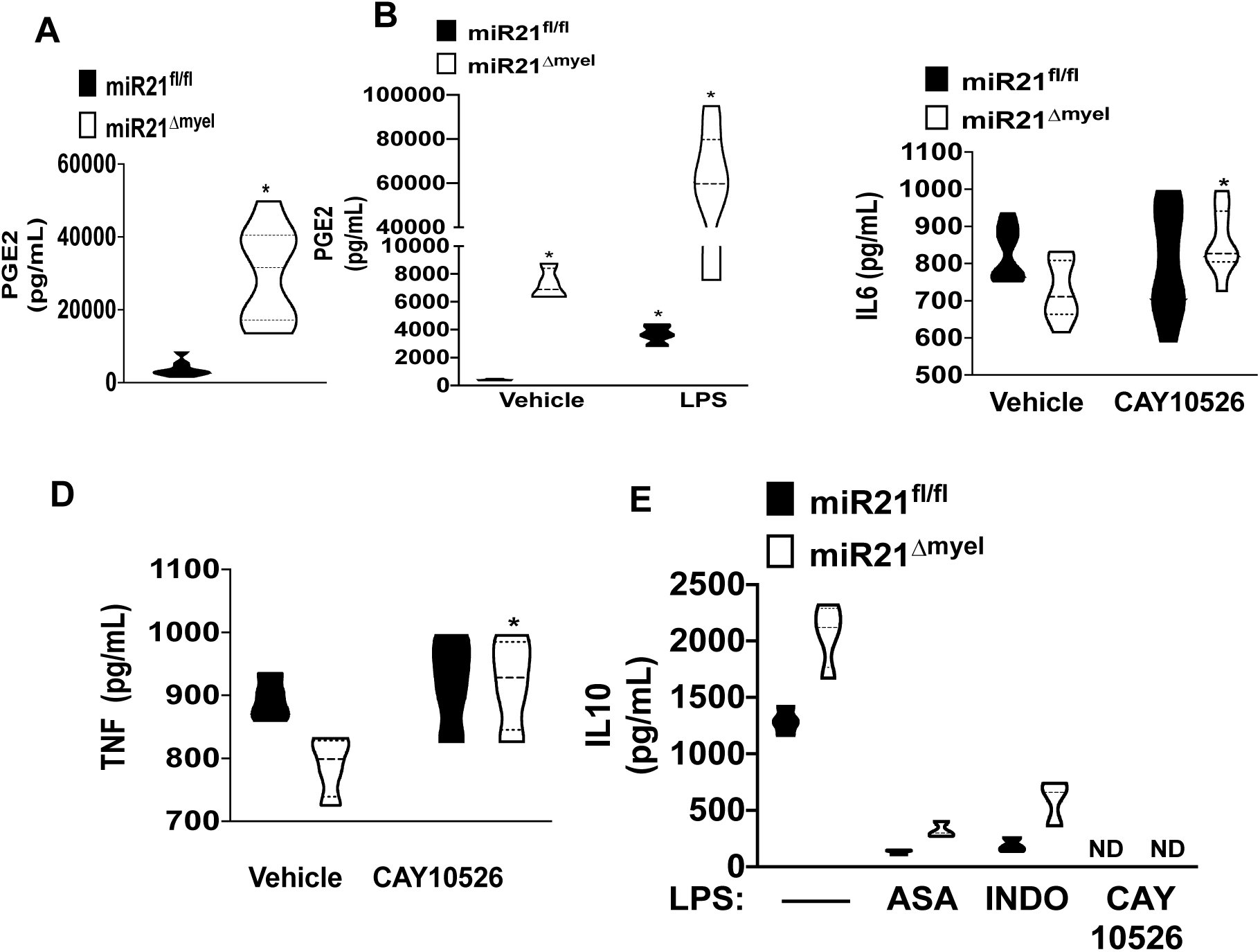
miR21 regulates PGE_2_/IL10 axis. **(A)** PGE_2_ levels measured in the serum of miR21_fl/fl_ and miR2_Δmyel_LPS-induced septic shock mice 12h after LPS i.p injection. Data are from *n* = 5 mice per group and were analyzed by *t-test* and Mann-Whitney *U* test. **(B)** PGE_2_ levels measured in the supernatant of miR21_fl/fl_ and miR21_Δmyel_ BMDMs challenged or not with LPS (10ng/mL) for 6h by ELISA. Data are from 3 independent experiments and were analyzed by one-way ANOVA, followed by Bonferroni correction. **(C)** The concentration of IL-6 and TNF-α measured in the supernatant of miR21_fl/fl_ and miR2_Δmyel_ BMDM LPS-challenged previously treated with CAY10526 (1 μM) or DMSO (0.1%) for 6h by ELISA. **(E)** Concentration of IL-10 in measured in the supernatant of miR21_fl/fl_ and miR2_Δmyel_ BMDM treated with aspirin (ASA, 100 μM), Indomethacin (IDO, 10 μM) and CAY10526 (CAY, 1 μM) one hour before challenge with LPS (10 ng/mL) for 6h by ELISA. **B-E** data are from at least 3 independent experiments and were analyzed by one-way ANOVA, followed by Bonferroni correction. Violin plots show the frequency distribution of the data \**P* < 0.05 compared *in vivo* by miR21_fl/fl_ mice or *in vitro* in LPS-challenged miR21_fl/fl_ BMDM (control) or unstimulated.

### Enhanced IL10 production drives *miR21*_*Δmye*l_ protective effects in LPS-challenged *miR21*_*Δmye*l_

IL10 is classically considered an anti-inflammatory cytokine(57), and more recently, Ip, et al. have shown that the inhibitory effect of IL10 is dependent on the inhibition of LPS-induced glucose uptake and glycolysis and increased oxidative phosphorylation (58). Here, we are studying whether miR21/dependent IL10 production is the primary driver of endotoxin resistance in *miR21*_*Δmye*l_. Initially, we treated *miR21*_*Δmye*l_ BMDMs with an IL10 blocking antibody, as we have previously shown (44), followed by the LPS challenge for 24 h. When we determined the abundance of TNFα and IL6 in the supernatant of *miR21*_*Δmye*l_ BMDM, blocking IL10 actions further increased the production/secretion of these cytokines, indicating that IL10 responsible for low production of inflammatory cytokines in *miR21*_*Δmye*l_ macrophages **(Figure 7A).** Next, we aimed to determine whether the blocking IL10 would render *miR21*_*Δmye*l_ mice susceptible to endotoxin. Our data clearly show that anti-IL10 treatment reflects in increased mortality to endotoxin in LPS-challenged *miR21*_*Δmye*l_ mice **(Figure 7B)**, as well as restored production of IL6, TNFα, and IL1b in the serum of miR21deficient mice **(Figure 7 C, D, and E)**. Importantly, blocking IL10 in vitro also restored glycolysis in BMDM from LPS-challenged *miR21*_*Δmye*l_ mice **(Figure 7F).** Together, these data show a heretofore unknown link between mir21/glycolysis/PGE_2_ and IL10 in macrophage inflammatory response and resistance to sepsis/endotoxemia in mice.

**Figure 7:**
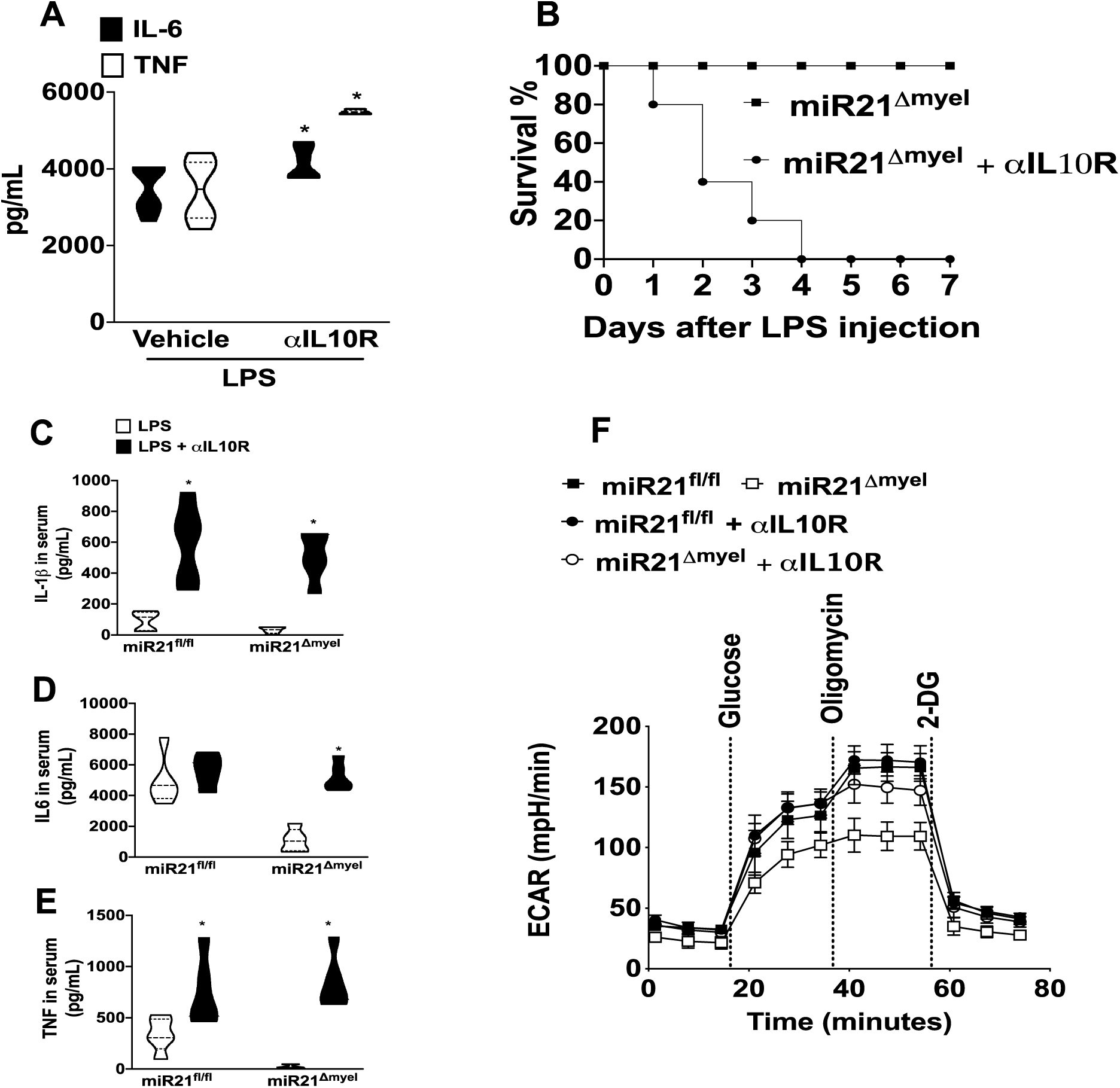
IL-10 receptor blockage prevents anti-inflammatory effects of miR21 deficiency. **(A)** Concentration of TNF-α and IL-6 levels in the supernatant of miR21_Δmyel_ BMDM treated with anti-10R (10 μg/mL) or PBS (Vehicle) 1-hour prior LPS challenged (10 ng/mL) for 12 hours by ELISA. Data are from *n* = 1-2 mice per group/experiment from at least 3 independent experiments and were analyzed by one-way ANOVA. **(B)** The survival rate of miR21_Δmyel_ mice challenged with a lethal dose of LPS (10 mg/kg) and treated with anti-10R (10 mg/kg) or PBS (Vehicle) 1 hour before and 24h after LPS i.p injection. Survival curve was analyzed for 7 days (n=5/7 mice per group) \**P* < 0.05 versus LPS/PBS treated miR21_Δmyel_ analyzed by log-rank (Mantel-Cox) test. **(C-E)** Concentration of IL-1β, IL-6, and TNF-α in the serum of miR21_fl/fl_ and miR21_Δmyel_ mice pretreated with anti-10R (10 mg/kg) or PBS (Vehicle) 1 hour before LPS i.p. injection (10 μg/kg). Cytokines were measured 24h later by ELISA. Data are from *n* = 5 mice per group and were analyzed by one-way ANOVA, followed by Bonferroni correction. **(F)** Real-time ECAR evaluation of glycolysis in miR21_fl/fl_ and miR21_Δmyel_ BMDMs treated with anti-10R (10 μg/mL) or PBS (Vehicle) 1-hour prior LPS challenged (10 ng/mL) for 12 hours by Seahorse XF glycolysis stress test. Data are from *n* = 1-2 mice per group/experiment from at least 3 independent experiments and were analyzed by one-way ANOVA. Violin plots show the frequency distribution of the data \**P* < 0.05 compared by miR21_fl/fl_ BMDM LPS (control) or unstimulated.

## Discussion

The outcome of sepsis is dictated by the impact of macrophage-induced production of inflammatory mediators along with the loss of structural cells and the release of danger-associated molecular patterns (59, 60). Macrophages are exquisite cells in their capacity to adapt to environmental clues; however, intracellular regulatory mechanisms must be in place to prevent macrophage hyperactivation (61). To avoid injury tissue, macrophage anti-inflammatory programs need to be adequately controlled at multiple levels, including transcriptional and posttranscriptional gene expression, post-receptor signaling events, and generation of metabolites that allow cells to adapt and respond to environmental cues (61, 62). Here we are employing complementary approaches to unveil a unique requirement of myeloid-specific miR21 in different arms of the macrophage biology resulting in the overwhelming inflammatory response, organ damage, and death in septic mice.

In summary, our findings show: 1) miR21 is highly expressed in macrophages and neutrophils after sepsis; 2) deletion of miR21 in myeloid cells enhances animal survival, decreases bacterial loads and cytokine storm and organ injury; 3) miR21 deficiency is accompanied by low glycolysis and expression of glycolytic enzymes; 4) miR21 inhibits the production of PGE_2_ that decreases the production of inflammatory cytokines and increases IL-10 abundance. 5) *In vivo* and *in vitro*, IL10R blockage restored cytokine production and glycolysis and enhanced mortality during endotoxin shock in *miR21*_*Δmye*l_.

miRNAs are fine-tuners of the induction, maintenance, and resolution of the inflammatory process, as well as antimicrobial effector functions (63). Although miRNAs control different aspects of the innate immune response, if cell-specific changes in the abundance of a single microRNA affect the outcome of sepsis, it remains to be fully determined. Several reports have shown that different microRNAs are involved in either the severity or resistance to sepsis (64, 65). Among the microRNAs, mir146a/, miR125a/b, miR155, and miR203 have all been involved in the pathogenesis of sepsis by directly controlling actions of TLRs, as well as inhibiting the activation of different transcription factors, such as NFκB, activating protein 1 (AP1) and cAMP-responsive element-binding (CREB) (66-68). In general, these well-studied microRNAs show anti-inflammatory properties and, in many ways, improve the outcome of sepsis in murine models (64, 69). On the contrary, few microRNAs show pro-inflammatory properties, but their targets and mechanisms remain elusive. Since a single microRNA could repress different targets, it is not uncommon to identify microRNAs that play opposite actions in a time-dependent manner (70-72). This is the case of miR21(33, 35).

miR21 is one of the most abundant microRNAs expressed during LPS challenge, cytokine (TNFα, IL1β, IL6, TGFβ, GM-CSF), and lipids (PGE_2_ and resolvin A1) stimulation (33). miR-21 is upregulated in septic patients and mice (**Figure 1** and (29). The role of miR21 in LPS-induced endotoxemia or sepsis is inconclusive. McClure *et al*., have shown that treatment of BALB/c mice with a miR21 antagomir enhances animal survival and decreases bacterial load after CLP (39). These findings also corroborated with Xue et al., that showed increased miR21 KO survival during sepsis by controlling IL1β-dependent inflammasome activation (40). Delivery of a miR21 antagomir was cardioprotective in septic mice by increasing the numbers of myeloid suppressor cells (38). On the other hand, Barnett et al. showed that miR21 KO mice are more susceptible to LPS-induced shock. Still, they did not identify any differences in animal mortality after CLP-induced sepsis (41). The reason for such discrepant data mentioned above might stem from different factors, 1) animal strains used, 2) the severity of sepsis, 3) source and quantity of LPS tested, 4) degree of gene silencing/inhibition using miR21 antagomir, as well as genetic and pharmacological approaches and global vs. cell-specific miR21 knockdown. Here, we are not only confirming the protective effect of miR21 deletion during sepsis and endotoxemia but also moving the field forward by dissecting the importance of the miR21 expression in myeloid cells (the critical cells involved in cytokine storm and organ damage during trauma) in the outcome of sepsis. Further studies are needed to determine whether the treatment of mice with miR21 antagomir or mimic after the induction of sepsis will also play a role in the outcome of sepsis.

There is no common scientific consensus about the exact mechanism of miR-21 in the induction or repression of the inflammatory program. Several manuscripts suggest miR-21 as an anti-inflammatory target (41, 73-75), while others suggested this miRNA as a crucial pro-inflammatory factor (36, 40, 76). We observed an overall decrease in IL6 and IL1b and an increase in IL10 abundance in septic miR21_Δmyel_ when compared to septic WT mice. Changes in cytokine production could be explained by the overall changes in metabolic changes in macrophages.

Immune cell metabolic reprogramming is a crucial part of the inflammatory response, and aerobic glycolysis is required for macrophage pro-inflammatory programs in myeloid cells (). However, aberrant aerobic glycolysis is associated with the development of detrimental systemic inflammation, and poor outcome in sepsis (21, 26, 77). We and others have shown that detrimental glycolysis is accompanied by increased HIF1a-dependent expression of glycolytic enzymes in both people and mice with sepsis (26, 54, 78).. Increased expression of individual glycolytic enzymes is also associated with enhanced mortality in sepsis (10, 54). Here, we observed decreased HIF-1α, GLUT-1, and HK-1, along with lower ECAR response and cytokine production. Also, we are showing that CD11b_+_ peritoneal cells and BALF cells from miR21_Δmyel_ septic mice when compared to septic WT, along with and increased animal survival. If decreased HI1Fα is the main responsible for overall lower glycolysis in miR21_Δmyel_ mice remains to be determined. Although lower glycolysis in miR21_-/-_ macrophages is not in agreement with the data from Hackett et al.,(79) showing increased expression of PFKM, ECAR, and production of inflammatory cytokines. Other reports have demonstrated that miR21 deficiency leads to decreased HIF1α-dependent glycolysis in different cancer cells (80, 81). The reason for the discrepancy between our findings and Hackett, et al., remains to be thoroughly dissected, but possible differences include the method used to differentiate BMDM (MCSF and GMCSF for 6 days) vs. 5% L929 supernatant throughout the experiments, and therefore, these cells are potentially different.

We and others have shown that PGE_2_ inhibits IL1β and TNFα, while enhances IL10 production in macrophages (82-84). Also, PGE_2_ is known to decrease glycolysis in different cell types(85, 86). Therefore, we speculated that PGE_2_ is a central hub for miR21 actions in macrophage glycolysis and cytokine production. Interestingly, we did observe increased PGE_2_ levels *in vivo* and *in vitro* in the absence of miR21 and inhibition of PGE_2_ synthesizing enzymes decreased IL10 and increased IL1β, IL6, and TNFα, suggesting that miR21 regulates the balance between PGE_2_ and IL10, controlling macrophage activation. That miR21 can target PGE_2_ synthesizing enzymes has been suggested in cancer models(87, 88). The enzyme that degrades PGE_2_ 15-PGHDS is targeted by miR21 in different cancer cell lines (87, 88). In our hands, we did not see any difference in 15-PGHDS expression in macrophages from WT and miR21_-/-_ macrophages. Whether miR21 mimics enhances 15-PGHDS expression remains to be determined. Importantly, we did not detect any effect of PGE_2_ inhibition, in ECAR levels, suggesting that a different mediator that is downstream to PGE2 actions is controlling glycolysis in miR21_-/-_ cells.

A plausible mechanism to explain changes in PGE2-mediated inhibition of cytokine production in miR21 deficient macrophage is the anti-inflammatory cytokine IL10 (57). The role of IL10 decreased glycolysis and cytokine production in miR21_-/-_ deficient mice and macrophages was confirmed using blocking IL10R. Sheedy et al. have shown that miR21 enhances IL10 abundance by directly controlling the expression of the IL10 inhibitor PDC4 (35). IL10 expression is regulated by different transcription factors, including STAT3 and CREB (89). CREB activation is governed by the actions of cyclic AMP-dependent protein kinase A (PKA)-mediated phosphorylation (55).

In summary, we are unveiling a heretofore unknown program by which a single microRNA regulates the actions of different classes of molecules, including lipid, protein, and metabolites that influences the outcome of an exaggerated inflammatory response triggered during sepsis.

