## Supplementary material for "Macrophage-derived miR-21 drives overwhelming glycolytic and inflammatory response during sepsis via repression of the PGE_2_/IL-10 axis": sup. fig 1-5

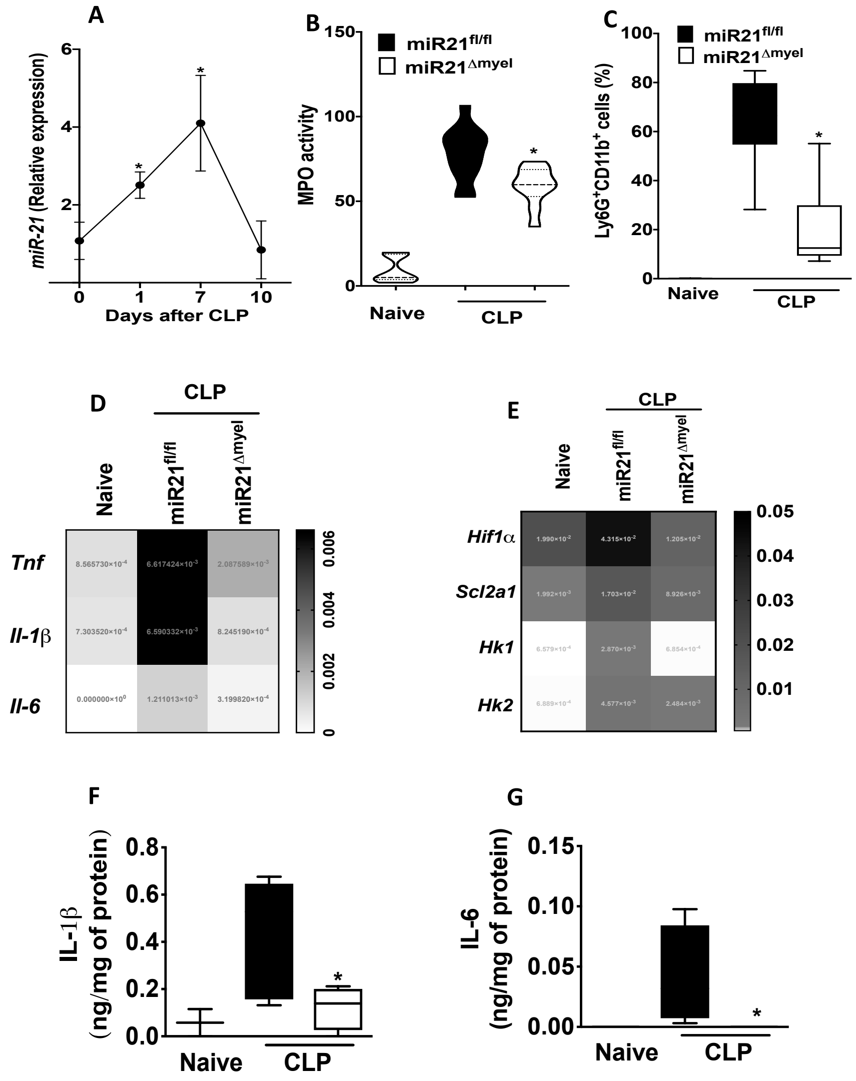


**Supplementary figure 1: miR-21 induces proinflammatory response in the lung of septic mice. (A)** miR-21 expression levels in the lung from WT septic mice 1, 7 and 10 days after CLP determined by qPCR. **(B)** MPO activity in the lung of miR21$^{fl/fl}$ and miR21$^{\Delta myel}$ naïve and septic mice 18h after CLP. **(C)** Percentage of Ly6G$^{+}$CD11b$^{+}$ cells in the lung of miR21$^{fl/fl}$ and miR21$^{\Delta myel}$ naïve and septic mice 18h after CLP by Flow Cytometer. **(D)** *Tnf, Il-1β, IL-6* mRNA expression in the cells of BALF miR21$^{fl/fl}$ and miR21$^{\Delta myel}$ naïve and septic mice 18h after CLP by qPCR. **(E)** *Hif-1α, Slc2a1, Hk1 and Hk2* mRNA expression in the cells of BALF from miR21$^{fl/fl}$ and miR21$^{\Delta myel}$ naïve and septic mice 18h after CLP by qPCR. **(F-G)** Concentration of IL-1β and IL-6 measured in the BALF exudate from miR21$^{fl/fl}$ and miR21$^{\Delta myel}$ naïve and septic mice 18h after CLP by ELISA. In all experiments, data are from $n$ = 4-7 mice per group and were analyzed by one-way ANOVA, followed by Bonferroni correction. Box-and-whiskers plots show Min. to Max and median. $P < 0.05$ compared to sham mice or miR21$^{fl/fl}$ septic (control) in all experiments.

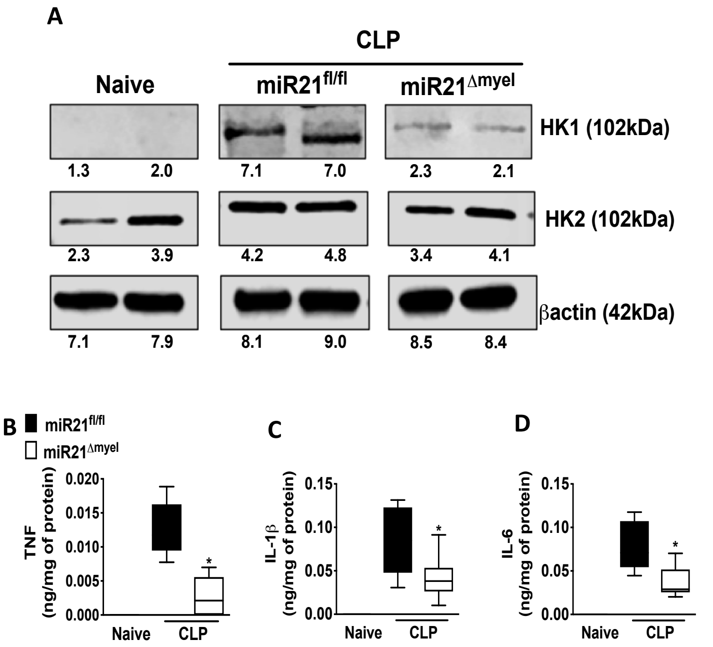


**Supplementary figure 2: miR21 regulates the expression of glycolytic enzymes and production of proinflammatory cytokines in peritoneal cells**. **(A)** HK1, HK2 and βActin expression of peritoneal cells from miR21$^{fl/fl}$ and miR21$^{\Delta myel}$ naïve and septic mice 18 h after CLP by Immunoblotting. **(B-D)** Concentration of TNF, IL-1β and IL-6 levels in the peritoneal exudate from miR21$^{fl/fl}$ and miR21$^{fl/fl}$ $\Delta^{myel}$ naïve and septic mice 18 h after CLP by ELISA. Data are from $n$ = 4-7 mice per group and were analyzed by one-way ANOVA, followed by Bonferroni correction. Box-and-whiskers plots show Min. to Max and median. $P$ < 0.05 compared to sham mice or miR21$^{fl/fl}$ septic (control) in all experiments.

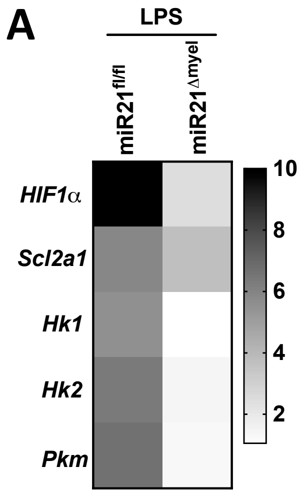


**Supplementary figure 3: miR21 is critical to LPS induced expression of glycolytic genes related in BMDM.** (**A**) *Hif1α, Scl2a1, Hk1, Hk2 and Pkm* mRNA expression in miR21$^{fl/fl}$ and miR21$^{fl/fl}$ $\Delta^{myel}$ BMDMs challenged with LPS (10ng/mL) for 12h determined by qPCR.

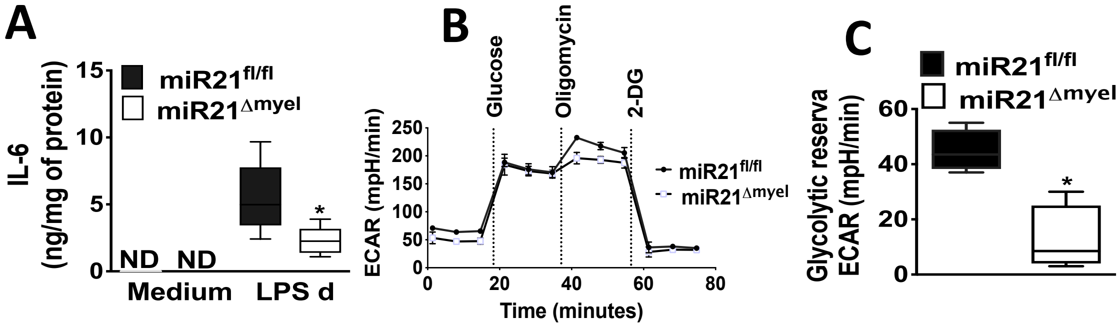


**Supplementary figure 4: miR-21 regulates IL-6 production and aerobic glycolysis in elicited peritoneal macrophage. (A)** Concentration of IL-6 measured in the supernatant of miR21$^{fl/fl}$ and miR21$^{\Delta myel}$ thioglycolate-elicited peritoneal macrophage challenged with LPS for 12h by ELISA. Data are from at least 3 independent experiments and were analyzed by one-way ANOVA. **(B,C)** Real time ECAR evaluation of glycolysis and glycolytic reserve of miR21$^{fl/fl}$ and miR21$^{\Delta myel}$ TEM LPS-challenged for 12h by Seahorse. Data are from least 3 independent experiments and were analyzed by *t* test and Mann-Whitney *U* test. Box-and-whiskers plots show Min. to Max and median. *$P$ < 0.05 compared by miR21$^{fl/fl}$ TEM LPS (control) or unstimulated.

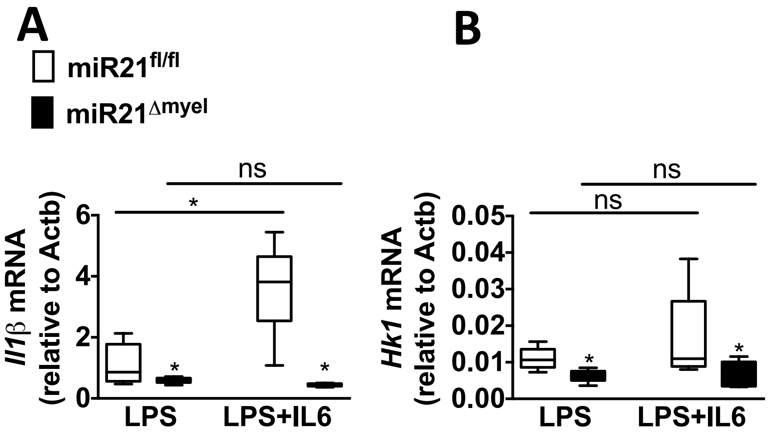


**Supplementary figure 5: IL-6 does not rescue the anti-inflammatory phenotype of miR21 deficient macrophages.** **(A,B)** *Il1b* and *Hk1* mRNA expression of miR21$^{fl/fl}$ and miR21$^{\Delta myel}$ BMDMs challenged with LPS (10 ng/mL) or LPS plus IL-6 (10 ng/mL) for 12 hours determined by qPCR. Data are from at least 3 independent experiments and were analyzed by one-way ANOVA. Box-and-whiskers plots show Min. to Max and median. *$P < 0.05$ compared by miR21$^{fl/fl}$ BMDM LPS (control).
